# Serological screening suggests single SARS-CoV-2 spillover events to cattle

**DOI:** 10.1101/2022.01.17.476608

**Authors:** Kerstin Wernike, Jens Böttcher, Silke Amelung, Kerstin Albrecht, Tanja Gärtner, Karsten Donat, Martin Beer

**Affiliations:** Friedrich-Loeffler-Institut, Greifswald – Insel Riems, Germany; Bavarian Animal Health Service, Poing, Germany; LUFA Nord-West, Oldenburg, Germany; State Institute for Consumer Protection of Saxony-Anhalt, Stendal, Germany; Thuringian Animal Diseases Fund, Animal Health Service, Jena, Germany

**Keywords:** SARS-CoV-2, COVID-19, animal, reservoir, cattle, ruminants, livestock, serology, epidemiology

## Abstract

Widespread human SARS-CoV-2 infections pose a constant risk for virus transmission to animals. Here, we serologically investigated 1000 cattle samples collected in late 2021 in Germany. Eleven sera tested antibody-positive, indicating that cattle may be occasionally infected by contact to SARS-CoV-2-positive keepers, but there is no indication of further spreading.

## Text

Since its first detection at the end of 2019, the betacoronavirus SARS-CoV-2 is keeping the world in suspense. This novel virus, which induces coronavirus disease 2019 (COVID-19) in humans, very rapidly spread around the world, thereby causing a massive global pandemic that resulted in more than five millions of deaths in less than two years of virus circulation *(1)*. Since the beginning of the pandemic, the role of livestock and wildlife species at the human-animal interface was discussed. A special focus was placed on the identification of susceptible species and potential intermediate or reservoir hosts. Under experimental conditions, various animal species could be infected with SARS-CoV-2, among them non-human primates, felines, canines, mustelids, white-tailed deer and several *Cricetidae* species, while e.g. poultry or swine are not susceptible *(2)*. For domestic ruminants such as cattle, sheep or goat a very low susceptibility was demonstrated following experimental inoculation, as only a small proportion of animals could be infected without animal to animal transmission *(3–5)*. Furthermore, 26 cattle exposed in the field to SARS-CoV-2 via contact to their infected keepers tested negative by RT-PCR *(6)*. However, given the very short time frame of only one to two days at which cattle test RT-PCR positive after experimental infection *(3,7)*, serological screenings could be more beneficial to identify previously infected animals, in order to estimate the rate of spill-over infections in the field.

Here, 1000 available samples of cattle kept in 83 holdings located in four German federal states (Bavaria, Lower Saxony, Saxony-Anhalt and Thuringia) were analyzed. The sampling dates were autumn 2021 and early winter 2021/22 when a massive wave of infections in the human population driven by the Delta variant of concern (VOC) occurred. Two to 20 randomly selected serum or plasma samples were analyzed per holding. Farm 31 was sampled twice, in between the animal owner was quarantined. Whether this quarantine was due to contact to an infected person or whether the owner himself tested SARS-CoV-2 positive is not known to the authors. All bovine samples were tested by an RBD-based multispecies ELISA performed as described previously *(8)*. During the initial test validation and during an experimental SARS-CoV-2 infection study in cattle, it could be shown that the ELISA does not cross-react with the bovine coronavirus (BCoV) *(3,8)*. Here, additional 100 cattle control samples randomly collected across Germany in 2016 were investigated and all of them tested negative.

Of the animals sampled in 2021, 11 cattle from nine farms tested positive by the RBD-ELISA, among them one animal kept in farm 31 and sampled after the quarantine of the owner (Figure 1). All but one (farm 8) positive ELISA results could be confirmed by an indirect immunofluorescence assay (iIFA) using Vero cells infected with the SARS-CoV-2 strain 2019_nCoV Muc-IMB-1 (multiplicity of infection of 0.1) as antigen matrix *(3)*. The titers ranged between 1/8 and 1/512, where the highest titer was measured in the seropositive animal from farm 31 (Table 1). To further confirm the reactivity towards SARS-CoV-2, the 11 samples that reacted positive in the RBD-ELISA were additionally tested by a surrogate virus neutralization test (cPass SARS-CoV-2 Surrogate Virus Neutralization Test (sVNT) Kit, GenScript, the Netherlands). This test allows for the detection of neutralizing antibodies by mimicking the interaction between SARS-CoV-2 and the host cell’s membrane receptor protein ACE2. It was reported to be highly specific but only moderately sensitive for animal samples, since it does not detect low antibody titers *(9)*. Four cattle samples scored also positive by the sVNT (farms 11, 31, 47 and 74; Table 1).

**Table 1.** Detailed information about the results of samples that tested positive by a multispecies SARS-CoV-2 RBD-based ELISA. iIFA = indirect immunofluorescence assay, sVNT = surrogate virus neutralization test (cPass SARS-CoV-2 Surrogate Virus Neutralization Test (sVNT) Kit, GenScript, the Netherlands; cut-off ≥ 30% positive and < 30% negative)

| Cattle farm/animal<br>number | RBD-ELISA<br>(corr. OD - status) | iIFA<br>(titer - status) | sVNT<br>(% inhibition - status) |
| --- | --- | --- | --- |
| 8/1 | **0.35 - positive** | <1/8 - negative | 6.1 - negative |
| 11/1 | **0.70 - positive** | **1/32 - positive** | **36.4 - positive** |
| 31/1 | **1.00 - positive** | **1/512 - positive** | **57.8 - positive** |
| 34/1 | **0.50 - positive** | **1/32 - positive** | 11.7 - negative |
| 42/1 | **0.65 - positive** | **1/16 - positive** | 5.5 - negative |
| 45/1 | **0.67 - positive** | **1/8 - positive** | 10.6 - negative |
| 45/2 | **0.33 - positive** | **1/16 - positive** | 9.0 - negative |
| 47/1 | **0.48 - positive** | **1/8 - positive** | **37.1 - positive** |
| 47/2 | **0.67 - positive** | **1/8 - positive** | 0.6 - negative |
| 72/1 | **0.52 - positive** | **1/16 - positive** | 4.7 - negative |
| 74/1 | **0.76 - positive** | **1/32 - positive** | **54.2 - positive** |

**Figure 1.**
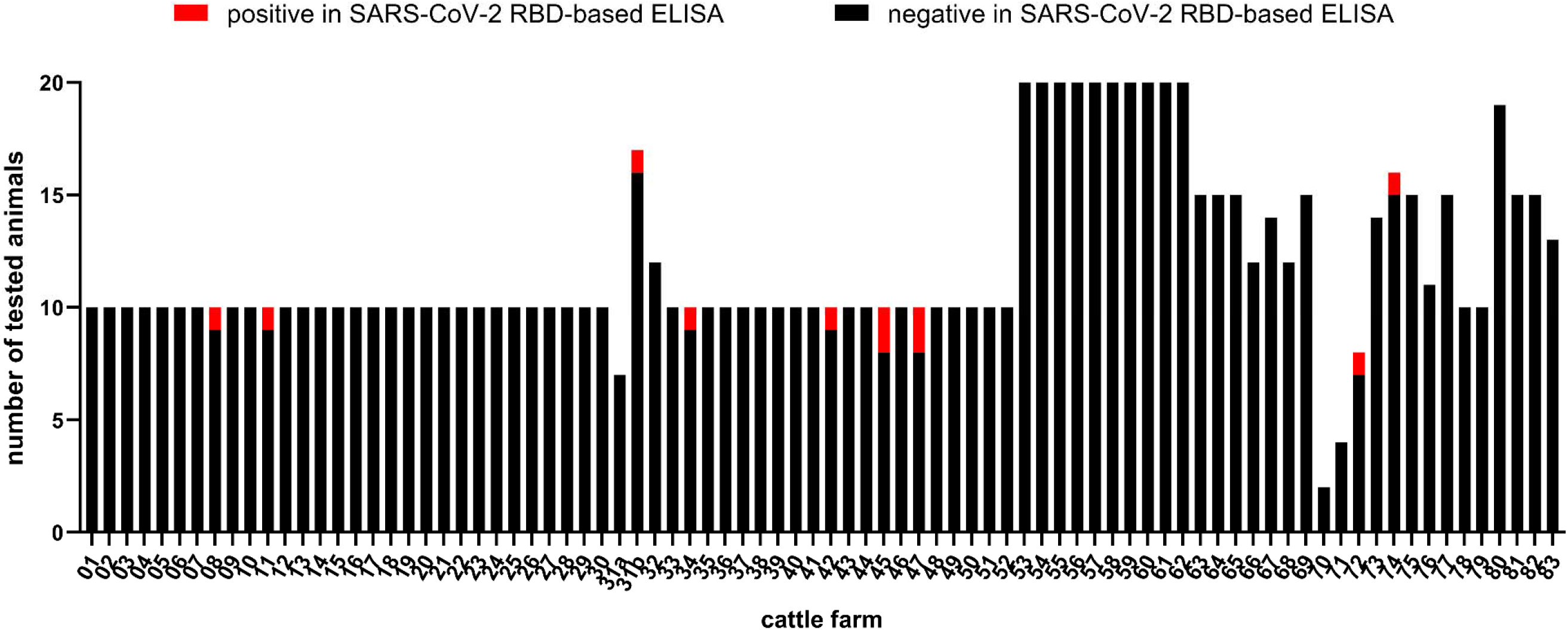
Number of cattle per farm tested for antibodies against SARS-CoV-2. Samples that reacted negative in the RBD-based ELISA are depicted in black and positive samples in red. Holding 31 was sampled twice (indicated as 31a and 31b), in between the animal owner was quarantined.

In conclusion, our findings of a low number of individual seropositive cattle in several farms demonstrate that cattle might be occasionally infected by contact to infected humans and seroconvert. However, in keeping with experimental infection studies *(3)*, intraspecies transmission seems likewise not to occur in the field. Nevertheless, cattle farms should be included in future monitoring programs, especially as another coronavirus, i.e. BCoV, is highly prevalent in the cattle population and a BCoV infection did not prevent a SARS-CoV-2 infection in a previous study *(3)*. Furthermore, we do not know the susceptibility of animal hosts for the new VOC Omicron.

Resulting double infections of individual animals could potentially lead to recombination between both viruses, a phenomenon well-described for other coronaviruses *(10)*. Although, the emergence is highly unlikely due to the low susceptibility of cattle for SARS-CoV-2, a conceivable chimera between SARS-CoV-2 and BCoV could represent an additionally threat. Hence, also ruminants should be included in outbreak investigations and regular screenings should be performed to exclude any spread of new variants in the livestock population.

## Acknowledgments

We thank Bianka Hillmann and Mareen Lange for excellent technical assistance. The study was supported by intramural funding of the German Federal Ministry of Food and Agriculture provided to the Friedrich-Loeffler-Institut.

## Ethical Statement

The serum samples represented superfluous material of routine diagnostic submissions taken by the responsible veterinarians in the context of the health monitoring of the respective cattle farm, no permissions were needed to collect these specimens.

